# Distance preserving dimension reduction with local-topology based scaling for improved classification of Biomedical data-sets

**DOI:** 10.1101/2019.12.27.889337

**Authors:** Karaj Khosla, Indra Prakash Jha, Vibhor Kumar

## Abstract

Dimension reduction is often used for several procedures of analysis of high dimensional biomedical data-sets such as classification or outlier detection. To improve performance of such data-mining steps, preserving both distance information and local topology among data-points could be more useful than giving priority to visualisation in low dimension. Therefore, we introduce topology preserving distance scaling (TPDS) to augment dimension reduction method meant to reproduce distance information in higher dimension. Our approach involves distance inflation to preserve local topology to avoid collapse during distance preservation based optimisation. Applying TPDS on diverse biomedical data-sets revealed that besides providing better visualisation than typical distance preserving methods, TPDS leads to better classification of data points in reduced dimension. For data-sets with outliers, the approach of TPDS also proves to be useful, even for purely distance-preserving method for achieving better convergence.

## Introduction

Dimension reduction of high dimensional data is an important problem in a wide variety of domains. Whether it is the field of genomics, proteomics or medical informatics, dimension reduction always pose a challenge to extract meaningful information in low dimension for visualisation, classification and other down-stream analysis. Traditionally three different kinds of approaches are used for dimension reduction which try to preserves one of the three factors: distances between data points, local topology or the overall information in the data. While principal component analysis is an information preserving method, multidimensional scaling (MDS) and Sammon mapping preserve distance [15]. Whereas, methods like t-SNE [10], and Locally-Linear embedding (LLE) [14] are nonlinear dimension-reduction techniques which aim to preserve local structure of data (Lee and Verleysen, 2007) [7]. These methods of dimension reduction have their own criteria and cost-function which they try to minimise. Such as Sammon mapping tries to minimise the squared error in scaled distances in high and low dimension normalised by the original distance in the high-dimensional space. One of the popular methods of dimension reduction, t-SNE aims to minimise cost function similar to SNE (stochastic neighbor embedding). SNE cost function [2] is based on Kullback-Leibler divergences between the conditional probabilities of distances based on Gaussian distribution. The SNE cost function does not allow collapse of similar data-points and it emphasizes on local distances which often leads to loss of information about the global topology. Methods like LLE and t-SNE reduce dimension of data-points to lower dimension coordinates which are dependent on properties of data. Some of the common dimension reduction approach proposed so far can also be catagorised as Manifold learning method [18]. Especially for image processing tasks such as segmentation, registration, tracking and recognition manifold learning has proved to be useful. A class of non-linear dimension reduction method like Self organising map (SOM) reduce the dimension of data to predefined coordinates (lattice). SOM is often used to make a initial prediction of manifold [4]. Even though SOM tends to provide visualisation of local neighborhood among groups of data-points, it is not able to properly represent local topology or global distance information.

Methods which are mainly designed for visualisation [3] [10] [17] often loose information which could be needed for other downstream analysis. On the other hand preserving global topology of distances, has its own importance in dimension reduction, especially for downstream analysis steps like clustering, phylogenetic analysis or regression. However, while optimisation of cost function of MDS, most often large distance dominate and cause collapse of data-points [1] which have some similarity among each-other but are distinct. Given recent trend in biological data-set, the collapse of data-points in lower dimension could lead loss of valuable and useful information. Such as single-cell gene-expression and proteomic data-sets are meant to highlight heterogeneity between cell-groups as well as among every cell so that gradient of cell-states can be studied [6] [19]. On similar trend, while studying cohort of individuals, we would like to visualise large difference between diseased and normal as well as exploit heterogeneity among disease cases for their stratification. Hence there is need of a method which can preserve pattern of larges distances as well the local topology due to heterogeneity. Such method could avoid artefact due to outlier large distances as well as avoid collapse of data-points and support downstream procedures like clustering and other machine learning based analysis steps.

Here, we describe an approach of dimension reduction such that both global and local topology could be preserved. We call our method as Topology preserving distance shrinkage (TPDS). TPDS uses the approach of distance shrinkage before using the method of non-metric MDS. The objective of TPDS is not only to improve visualisation like t-SNE but represent data in low dimension such that other procedures like classification could be made more efficient. Here we first provide the description about TPDS then explain the cost function involved in learning its parameters. Then in result section, we have shown results for 4 data-sets. In addition to visualisation we have used other classification in reduced dimension to evaluate the output of dimension reduction methods.

## Methods

We follow the approach of distance scaling so that even if distance preservation based method is used local topology remains intact. Our approach is inspired by Einstein’s field equation which suggest relativity and space-time warping [9]. Hence even if light travels in straight line, the warping of space-time bends it’s path.

Similarly in our approach MDS method tries to preserve the distances, however due to warping of distances among data-points, the local topology could also be preserved. However for this purpose some prior guess of manifold and local topology is needed. Therefore, we first use self organising map (SOM) to get a rough estimate of groups and initial prediction of manifold. The grouping of data-points by SOM need not to be necessarily correct but it provides an estimate of neighborhood to be utilised by TPDS. Moreover if the number of features are too large, one can first perform singular value decomposition of data and use singular vectors with SOM. After getting a rough estimate of manifold and neighborhood, we scale distance between the data-points using their belonging-ness to different groups and distance between group means. The scaling function used by TPDS consists of an attractive and a repulsive component. For this purpose we define *m*_*ij*_ as likelihood of proximity between two data-points *i* and *j* belonging to two different SOM clusters *m* and *k* as

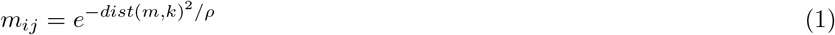

Here *dist*(*m, k*) is the normalised distance between centers for cluster *m* and *k*, and *ρ* is spread factor just like variance in Gaussian distribution. The *dist*(*m, k*) is normalised by division with mean of all distances between cluster centers. If data points *i* and *j* belong to same SOM cluster the value of *m*_*ij*_ is calculated using distance *d*_*ij*_ between them

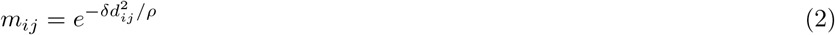

Notice that variable *ρ* is same as in equation(1) and *δ* is a multiplying factor less than 1. We calculate the scaling factor for distance between data-points *x*_*i*_ and *x*_*j*_ using the formula.

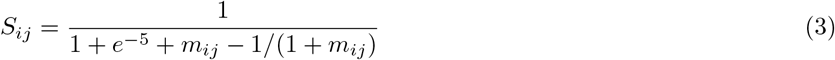

The purpose of *δ* is to inflate local distances wherever the data-points lie in same cluster so that we can avoid their collapse. After calculating scaling factor, the distance between data-points are scaled using the formula

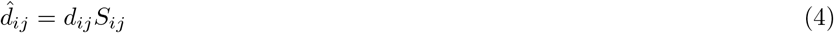

The matrix of scaled distances *d̂*_*ij*_ is then used with non-metric MDS method to reduce the dimension. Notice that when likelihood of proximity *m*_*ij*_ is low the scaling factor *S*_*ij*_ increase and inflate the distance, hence it behaves like a repulsive force. However when likelihood *m*_*ij*_ is high and reaches closer to 1, the value of *S*_*ij*_ decrease to create an attractive force to shrink the distance *d*_*ij*_. Thus we inflate and shrink distances according the likelihood of proximity between data-points or their centers of clusters. Also notice that likelihood is calculated using two different ways, depending on the condition whether two data-points belong to same SOM cluster or not. If data-point belong to same SOM cluster, likelihood of their proximity is decreased so that the distances between them are inflated slightly. Such inflation of distances of data-points belonging to same cluster causes preservation of local topology, while optimising for distance preservation based dimension reduction.

### Adjusting spread factor for balance between distance-stress and local topology

The spread factor *ρ* in equation (1) indirectly control the level of scaling of distances. If the value of *ρ* is low, the likelihood of proximity for larger distances become negligible and local topology neighborhood is also small. Whereas if *ρ* is high, larger distances are not ignored while scaling, which would lead to preservation of global structure of distances but may suppress local topology information. Hence optimisation for *ρ* is an important step for TPDS. For optimisation of *ρ* we use combination of two well-known costs namely distance-stress and a local-topology preserving strain. TPDS chooses that value of spread factor *ρ* which minimise the cost function defined as

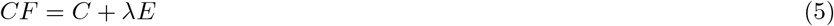

Where *λ* is lagrange multiplier. *C* is distance-based stress which measure the preservation of global distances pattern. *E* is a local-topology preserving constraint. TPDS uses default value of *λ* = 1, however different value of *λ* could also be used. Notice that we do not differentiate the cost function mentioned in equation (10) to learn *ρ* but choose the best value in a given range. It is just meant to choose the correct scaled distance matrix to be provided to the non-metric MDS method for dimension reduction. Thus TPDS also provides flexibility in choosing the level of preservation of local topology through *λ*.

Distance-based stress is the cost function used by MDS, which can be defined as sum of differences among distances in high dimension and reduced dimension

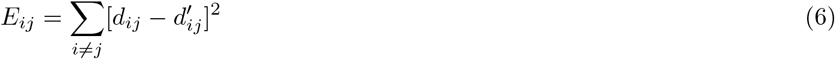

Here *d*_*ij*_ is distance between two data-points *x*_*i*_ and *x*_*j*_ in higher dimension space and 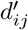 is distance in reduced dimension. The cost represented in eq (5) is often called at distance stress, which we term here as MDS-cost.

The strain of local topology preservation could be formulated using different approaches based on hard KNN or smooth neighborhood preservation [18] [13]. To test our model, we have used the cost function associated with symmetric SNE. As proposed by Lauren et al. [10] symmetric-SNE is a modification of SNE based approach. The symmetric-SNE cost function consists of single Kullback-Leibler divergence between a joint probability distribution, P, in the high-dimensional space and a joint probability distribution, Q, in the low-dimensional space

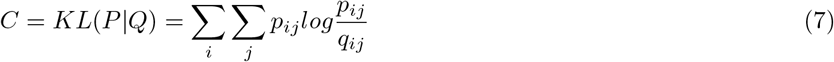

where *p*_*ii*_ and *q*_*ii*_ is set to zero. Here it is called as symmetric SNE because *p*_*ij*_ = *p*_*ji*_ and *q*_*ij*_ = *q*_*ji*_ as these pair wise similarities are defined as

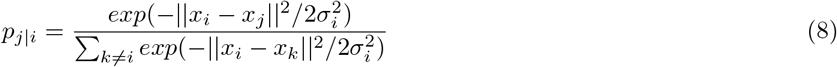

and

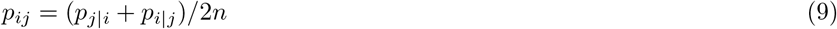

in higher dimension. Here *n* is the number of data-points. We keep the value of *σ_i_* at perplexity=15 [10].

In lower dimension the likelihood of distances is determined as

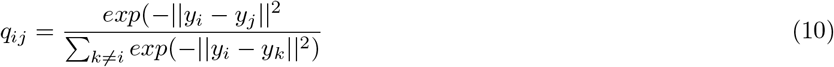

Here *y*_*i*_ and *y*_*j*_ represent coordinates of data-points in lower dimension.

## Results

In order to evaluate TPDS we used data-sets where both classification and visualisation is needed for making useful inference. The first data-set used for evaluation was generated by Naranjo et al. [12] and it consists of feature extracted from speech recording of normal individuals and Parkinon’s disease patients. For every patient or individual there are 4 replicate of data points which have 44 features. In the original manuscript, authors have used supervised classification approach to classify the normal and Parkison’s cases. We applied our unsupervised method of reducing the dimension. We also compared the output of our method with other unsupervised dimension reduction methods such as t-SNE, non-metric MDS and Sammon Mapping. As shown in the Figure 1 for Naranjo et al data-set [12] the visualisation itself shows two separate clusters for normal and Parkinson’s disease. Sammon mapping and MDS have similar separability of two classes like t-SNE. However TPDS revealed clear separability between Parkinson’s and normal cases and had lower MDS-cost than t-SNE function compared to Sammon mapping and non-metric MDS methods for dimension reduction. Further, we performed k-mean clustering (k = 2) on the output from TPDS and other tested methods. We calculated clustering purity using adjusted Rand index (ARI) and Normalized mutual information (NMI). K-mean clustering on output from TPDS had highest purity as the ARI and NMI scores were almost 1.7-2 times greater than other method used (tSNE, non-metric MDS and Sammon mapping). We also checked clustering purity after applying dbscan to cluster data-points in low-dimension. Even with dbscan based classification TPDS provided best clustering purity among tested methods (see table 1).

**Table 1.** Result of clustering purity after applying dbscan.

| Data-set | TPDS | t-SNE | non-metric MDS | Sammon mapping |
| --- | --- | --- | --- | --- |
| Parkinson's Naranjo et al. | ARI:0.43<br>NMI:0.33 | ARI: 0<br>NMI: 0 | ARI: 0.0<br>NMI:0.0 | ARI: 0.0<br>NMI: 0.0 |
| Mouse Protein Higuera et al. | ARI:0.133<br>NMI:0.23 | ARI: 0<br>NMI: 0 | ARI: 0.004<br>NMI:0.078 | ARI: 0.002<br>NMI: 0.068 |
| SCADI Zarachi et al. | ARI:0.288<br>NMI:0.355 | ARI: 0<br>NMI: 0 | ARI: 0.058<br>NMI:0.060 | ARI: 0.002<br>NMI: 0.068 |
| single cell Expression Li et al. | ARI:0.027<br>NMI:0.068 | ARI: 0.0022<br>NMI: 0.024 | ARI: 0<br>NMI:0 | ARI: 0<br>NMI: 0 |

**Figure 1.**
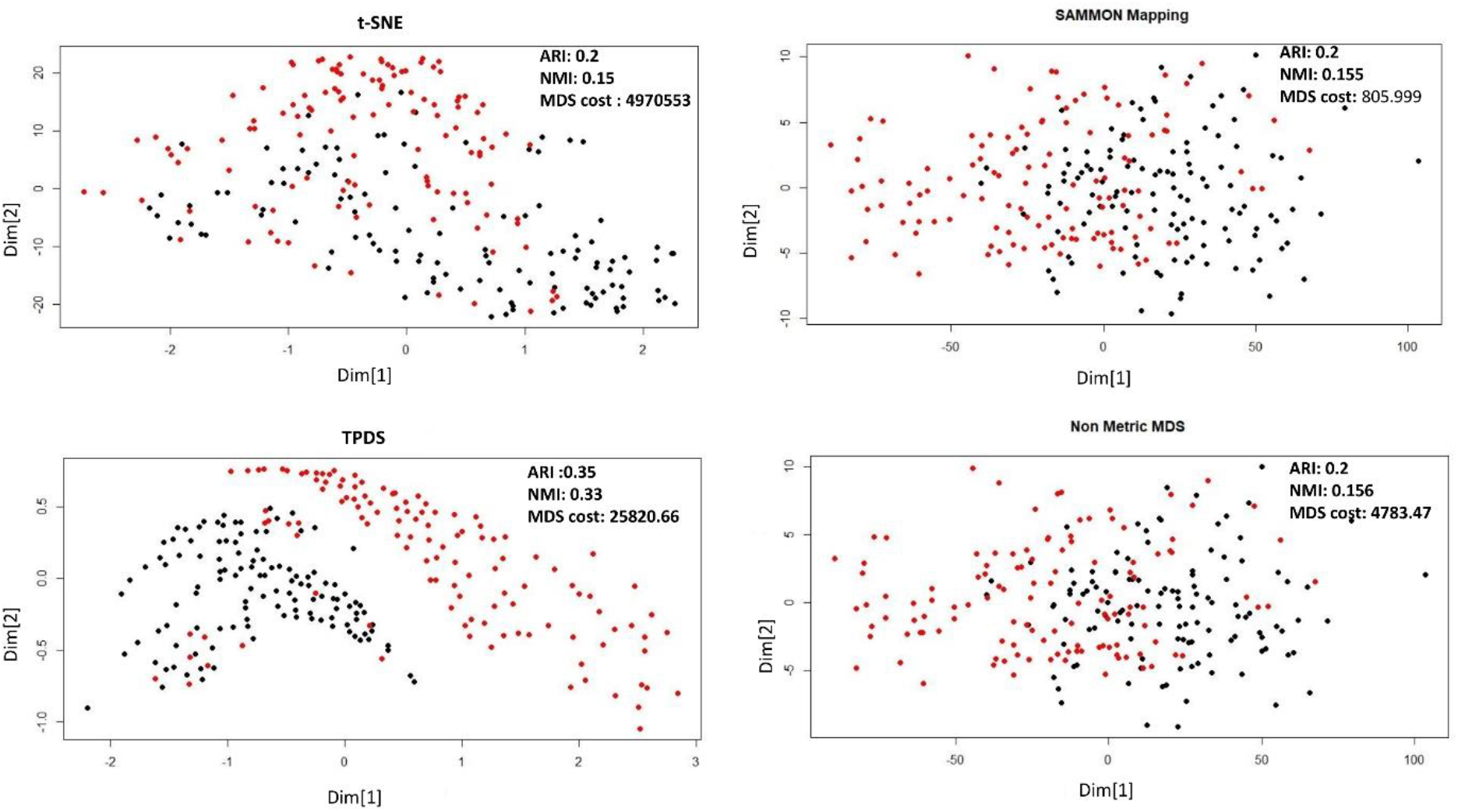
Dimension reduction of Parkinson’s data-set by Naranjo et al. [12] using four different method. The distance stress based cost is represent termed as MDS-cost

**Figure 2.**
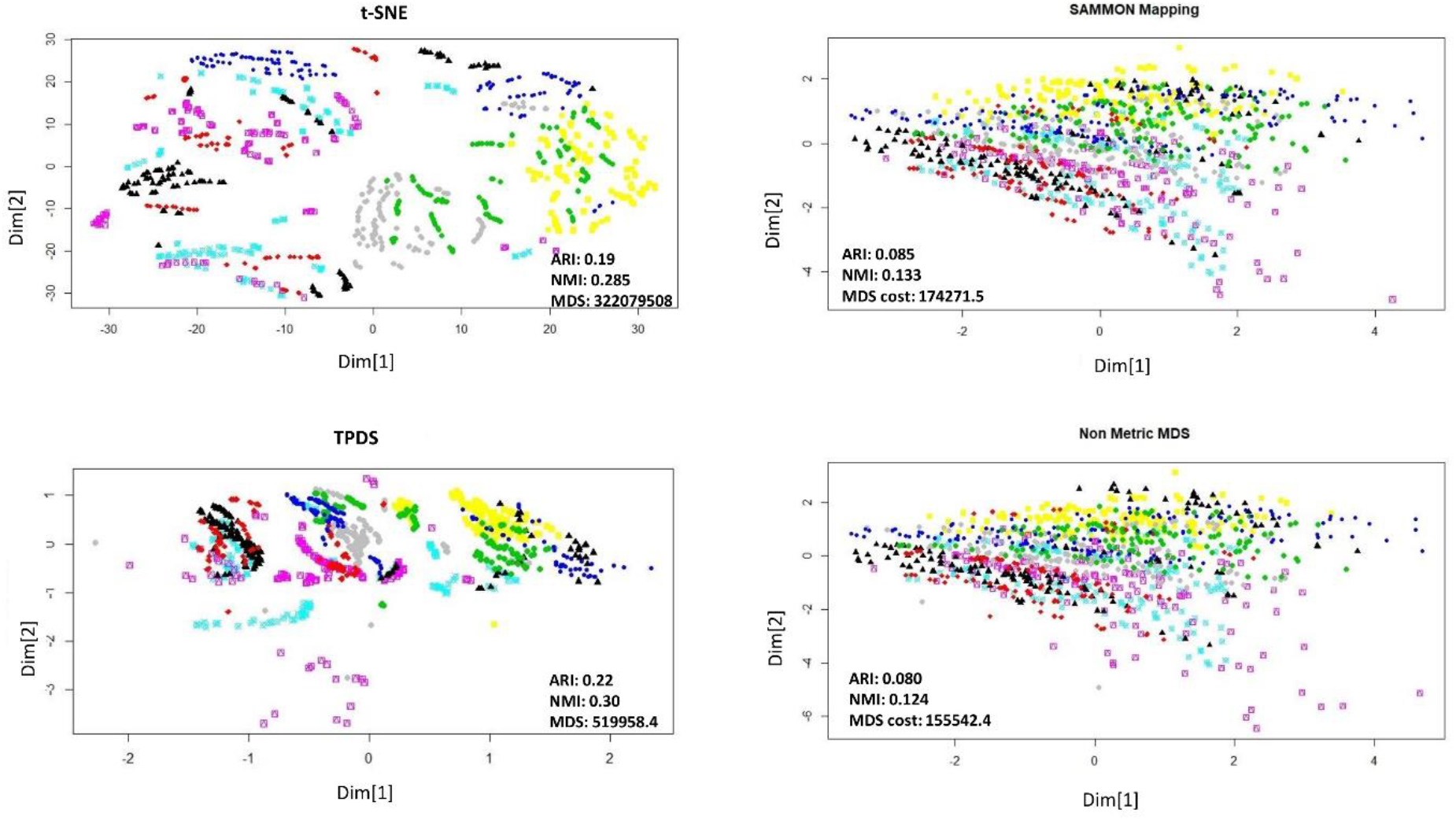
Dimension reduction of mouse protein data-set. Visualisation of reduction to 2 dimension. The distance stress based cost is represented as MDS-cost

**Figure 3.**
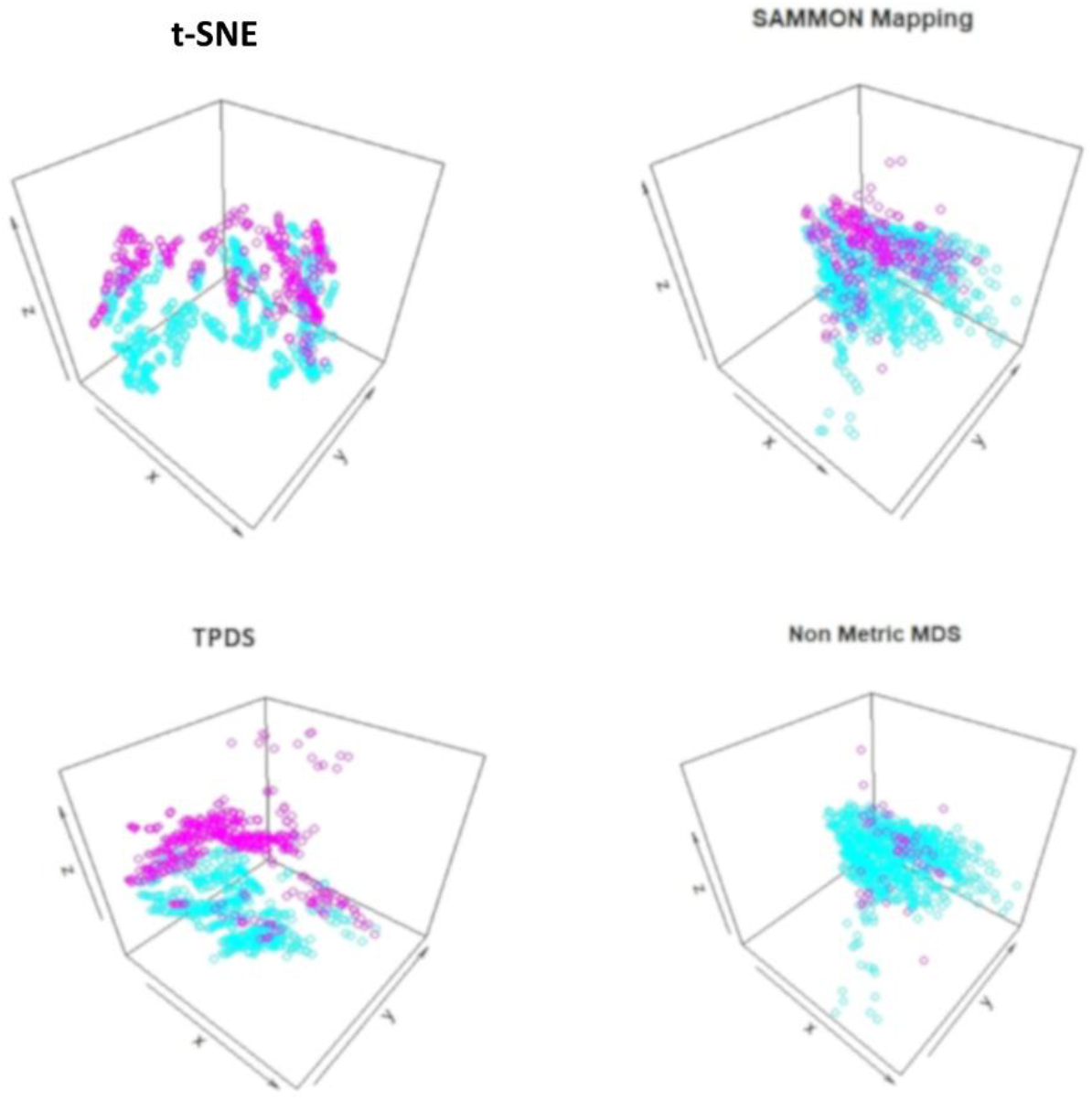
Visualisation of mouse protein data-set after reduction to 3 dimension. The trisomic and normal mice data-points are shown with different colors

**Figure 4.**
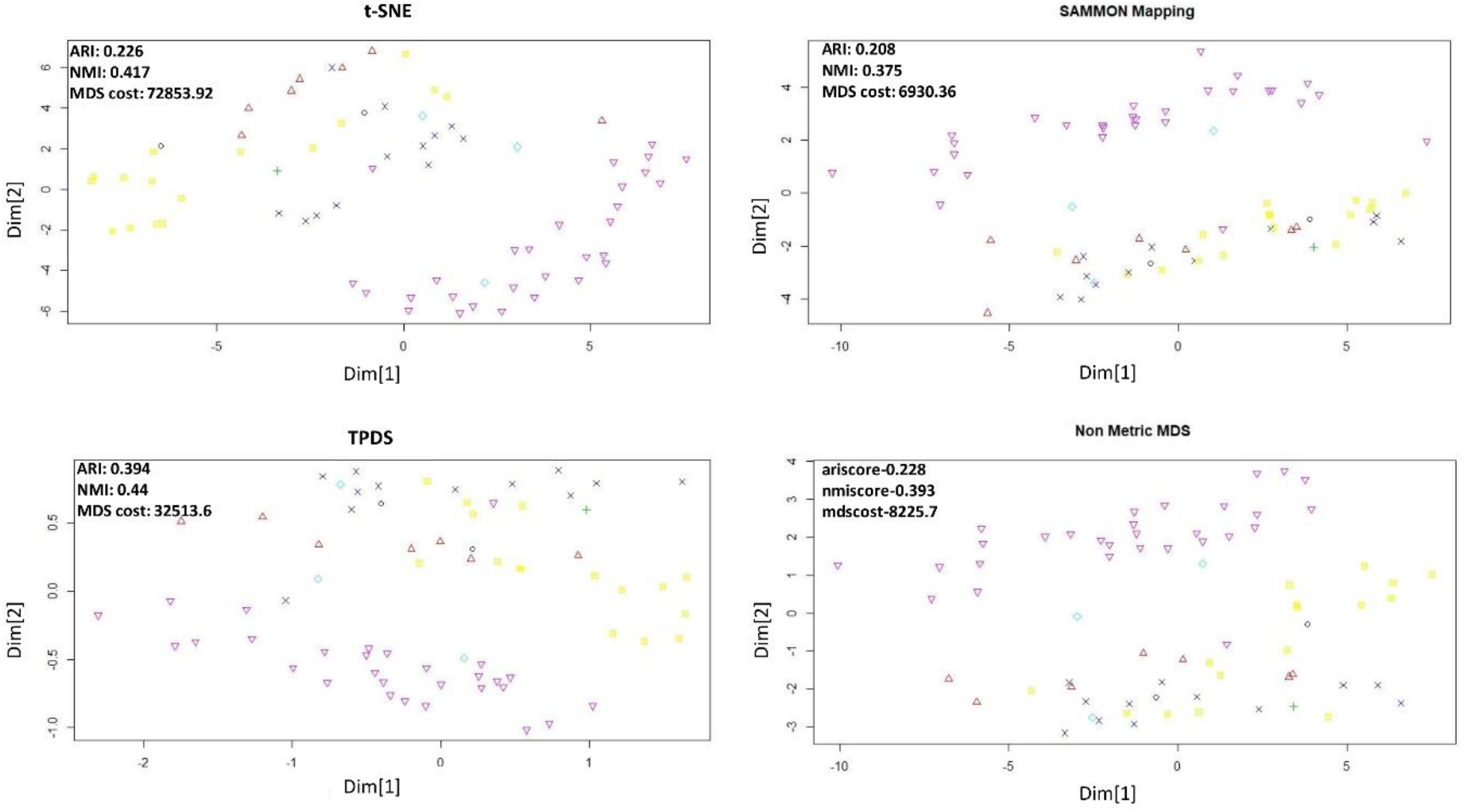
Dimension reduction of SCADI data-set.The adjusted Rand Index and Normalized mutual information (NMI) were calculated after performing k-mean clustering using k=7.

The second data-sets we used for evaluation, consists of the expression levels of 77 proteins/protein modifications from the the nuclear fraction of cortex of mouse [5]. This data-set of protein expression, was generated using 38 control mice and 34 trisomic mice with Down syndrome. There are 15 replicates per sample/mouse such that control mice, there are 38×15, or 570 measurements, and for trisomic mice, there are 34×15, or 510 measurements. Authors recommend to consider each replicate as a separate sample. The mice themselves belonged to 8 different groups depending on genotype, behavior and treatment. Using genotype mice could grouped as control or trisomic [5]. According to behavior, the mice could be grouped as stimulated to learn (context-shock) and not trained to learn (shock-context). Similarly mice could be grouped as treated with drug and others not. First, we visualised data-points with reduced dimension, using different colors for 8 different groups. Visually, it was non trivial to judge about which method performed better. The t-SNE method avoided crowding but spread out samples too much to cause mixing between different groups. With TPDS there seem to be crowding among some samples but for majority of samples co-localisation was according to group. On the other hand, Sammon mapping and non metric MDS caused data-points to be localised as highly overlapping strata according to 8 groups. Using K-mean clustering and calculating purity of classification provided higher NMI and ARI score for TPDS compared to other tested methods. We confirmed the improvement provided by TPDS in clustering using dbscan (see table 1). However, the mixing of classes hinted towards unknown co-variates. Therefore we again used TPDS to represent the high dimensional protein expression data-set using 3 dimensions. When data was reduced to three dimension, the 3D scatter plot of TPDS could show clear separability between trisomic and normal mice. Whereas other methods could not provide comparable separability like TPDS. Overall with mouse cortex protein expression data-set, we realised that TPDS has better potential to reduce dimension to align similar class data-points together and provide separability between non-similar groups.

The third data-set we used for evaluation is SCADI data-set [20] which was also downloaded from UCI database. The SCADI data-set contains 206 attributes of 70 children with physical and motor disability. The data-points are divided into 7 classes based on behaviour of children [20]. Applying our approach and other 3 methods revealed clear separability for few classes in reduced dimension. However, evaluating the separability after dimension reduction of SCADI data-set, revealed that classification of TPDS output resulted in substantially higher clustering purity than other tested methods (see Figure-1 and Table-1).

Finally we used single cell expression data to evaluate the potential of TPDS. The single cell expression data-set used here was published by Li et al. [8]. This data-set consists of read-count of more than 57242 genes as feature and 562 samples (cells). For dimension reduction of such data-sets with such large features, principal components are often used, especially in case of t-SNE. Therefore we provided loading on top 30 principal components to all the methods. The value of perplexity parameter for t-SNE was set to 4. The single-cell data-sets used here had labels for each cell and they could be categorised to 7 cell-types which we took as classes. The visualisation of t-SNE and TPDS outputs showed distinguishable loci for cells belonging to different types (classes). In terms of clustering purity using k-means and dbscan, TPDS had better performance than other methods. Surprisingly, the distance stress cost (MDS-cost) for TPDS was lower than other tested methods despite decent visualisation through preservation of local topology among cells. It hints about possible convergence issue of Sammon mapping and non-metric MDS method due to outlier cells. In Figure 5, it is clearly visible that cells (represented by red color) are outliers which caused have collapse of locations of other cell types with Sammon mapping and non-metric MDS. Notice that, even if TPDS uses nonmetric MDS method as last step, the collapse of data-points do not happen due to distance scaling to preserve local topology. Thus TPDS seem to have avoided local minima by providing less weightage to large distances of outlier data-point.

**Figure 5.**
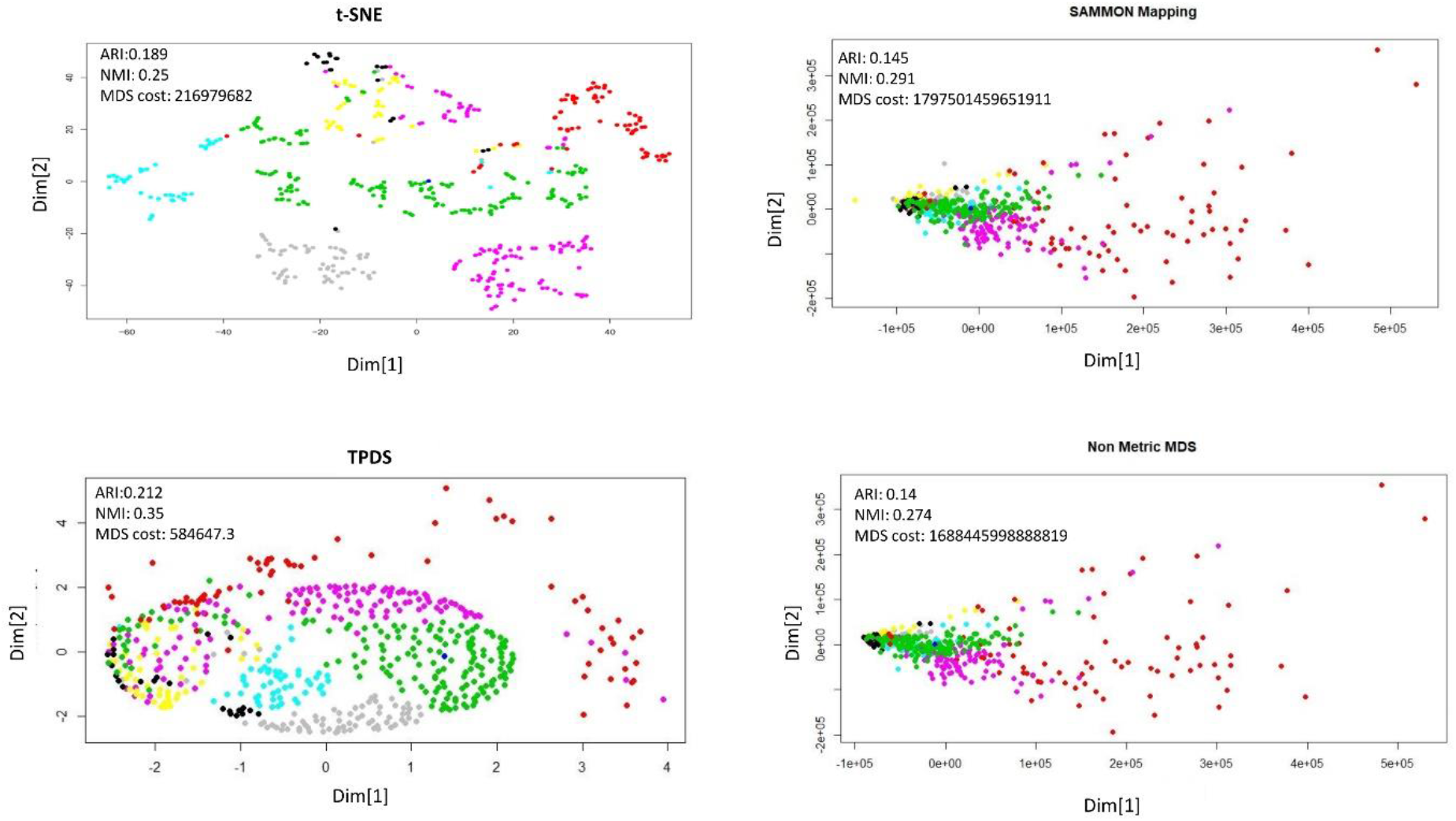
Dimension reduction of single cell expression data. The 7 types of cells are shown with different colors.The adjusted Rand Index (ARI) and Normalized mutual information (NMI) were calculated after k-mean clustering (k=7). The outlier cells (shown in red color) seem to have different effect in different dimension reduction methods

## Discussion

Various procedures of analysis of large data-sets such as classification, regression and anomaly detection can be improved using dimension reduction techniques. Given the diversity of Biomedical data-set, there could be multiple factors which influence classification. Hence techniques meant only for visualisation just as t-SNE may not be optimal for improving performance of analysis and classification for most of the data-sets. Preserving distance during dimension reduction could have it’s own advantage in terms of providing clear separability among dissimilar data-points. The success of TPDS in achieving low MDS-cost (distance stress), decent visualisation and providing the best clustering purity for tested data-sets hint that having a balance between local-topology and distance preservation could be useful for other analysis procedures.

The approach of TPDS to scale the distances to preserve some information about local topology can also help in better convergence of MDS like methods. It could be explained as such: the larger distances among data-points could act like outliers and cause hurdle in convergence. The smaller distances among data-points with in same group could be large in number, so giving them more weightage lead to better reduction of overall MDS-cost. In this process, some information about local topology could also be preserved for better visualisation. One important feature in results of TPDS is that besides separability among data-points of different classes they have tighter co-localisation among data-points of same classes, that too without collapse of their location. Such result caused substantially higher classification purity for TPDS results by dbscan which exploit such tight co-localisation for density based clustering. Hence TPDS could be used as an alternative dimension reduction method to support such density based clustering approach.

Here we used SOM to have preliminary estimation of manifold. Other methods generally use KNN or a modified version of it [11], [16] [14] for the same purpose. SOM provides neighborhood information about cluster of data-points, hence in comparison to KNN, SOM results could be less influenced by noise because of averaging effect. With TPDS one need to provide a guess about number of SOM clusters, which could have some effect on visualisation. However the number of SOM clusters would have less effect on the performance of TPDS for improving downstream analysis steps like classification and regression. Other methods like t-SNE and Sammon mapping are also dependent on parameters. t-SNE is dependent on perplexity parameter, whereas Sammon mapping used lambda as initial value of the step size during optimisation. t-SNE based dimension reduction for single cell expression data could be better at other values of perplexity, however our result shows that TPDS can also provide comparable results at optimal parameters. Thus TPDS is not an exception in being dependent on external parameters. In future we intend to make the guess about optimal number of SOM clusters for visualisation to be fully automatic. Overall, we have presented here a technique to utilise method for distance preservation based dimension reduction.

## Abbreviations

SNE: Stochastic Nieghorhood embeding
MDS: Multi-deimensional scaling
LLE: Locally-Linear Embedding
SOM: self Organing Map
KNN: K-nearest Neighbor
TPDS: topology preserving distance scaling

